# Berberine impairs spermatogenesis in mice by modulating ornithine metabolism via gut microbiota regulation

**DOI:** 10.1101/2023.10.30.564855

**Authors:** Wei Qu, Yumin Xu, Junhao Lei, Jing Yang, Hanqing Shi, Junli Wang, Xinnai Yu, Jiemin Chen, Binyi Wang, Yan Han, Mengcheng Luo, Rong Liu

## Abstract

Berberine (BBR) is used to treat diarrhea clinically, its reproductive toxicity, however, is poorly documented. This study aims to investigate the impact of BBR on the male reproductive system. Gradient doses of BBR were administered orally to experimental mice for consecutive 14 days. The gut microbiota, sperm concentration of cauda epididymis, and serum testosterone levels were measured after the last dose for assessing the effects of BBR. Moreover, the metabolome and transcriptome of the mice and microbiota were also investigated. Intragastric BBR administration resulted in a significant decrease in serum testosterone levels and epididymal sperm concentration in mice, which was attributed to a dramatic decrease of Muribaculaceae abundance in the gut microbiota of mice. Both fecal microbiota transplantation (FMT) and polyethylene glycol (PEG) treatment experiments also demonstrated that Muribaculaceae is necessary for spermatogenesis. Metabolomic analysis revealed that BBR affected the arginine and proline metabolism pathways, of which ornithine levels were downregulated after BBR administration. Intragastric administration of *M.intestinale* and its metabolite ornithine to BBR-treated mice achieved a recovery of sperm concentration and testosterone levels. RNA sequencing of testes showed the genes related to the LDLR-mediated cholesterol-synthesis testosterone pathway were downregulated after BBR administration. The levels of testosterone increased and *Ldlr* gene became more transcriptionally active in TM3 cells cultured in media supplemented with ornithine. This study for the first time revealed an association between BBR-induced gut Muribaculaceae dysbiosis and defects in spermatogenesis via ornithine metabolism, which provided a candidate and strategy for the treatment of infertility caused by a decreased serum testosterone level-induced by gut microbiota dysbiosis.

## Introduction

The incidence of infertility in couples of reproductive age is as high as 15%, of which male infertility accounts for approximately 50% [1]. Over the past 40 years, the seminal sperm concentration of adult males has decreased by over 50% in Western countries, which results from multiple factors, such as unhealthy diet and lifestyle, exposure to toxic chemicals, and irrational administration of medications [2]. Traditional Chinese medicine (TCM) has been widely used in clinical practice for disease prevention and treatment, but accumulating evidence shows that reproductive dysfunction could be caused by some of them. Triptonide, a traditional Chinese medicine-derived drug used for the treatment of immune-related diseases, was reported to induce sperm malformation and decreased motility without expectation [3].

Gossypol, a toxic crystalline compound present in cotton-seed oil, has antiviral and antimicrobial activities, but it has also been shown to repress the intercellular communication vai gap junction among Sertoli cells and steroidogenesis of Leydig cells in mouse testes [4, 5]. Aristolochic acid has been used to treat tumors, rheumatism, and arthritis; however, it also induces testicular injury through the inhibition of amino acid and glucose metabolism in male mice [6]. In recent years, berberine (BBR) has been used to treating diarrhea [7], inflammatory diseases [8], and colon cancer [9]. However, whether BBR has a definite effect on the male reproductive system remains to be elucidated.

Gut microbiota, as a potential organ, is involved in many important physiological processes of the host, including regulation immunity, maintaining metabolic homeostasis, maintaining cognitive functions and so on [10, 11]. Numerous herbal ingredients have low bioavailability and function by regulating gut microbiota. Saponins was reported to facilitate the growth of beneficial bacteria and to supress cachexia-like symptoms in mice [12]. Rhein, one of the main components of rhubarb, increased the abundance of *Lactobacillus* and alleviated uric acid-induced chronic colitis [13]. Piperine improved Parkinson’s disease by down-regulating the high abundance of *E. faecalis* [14]. BBR also has low bioavailability, so the gut microbiota is likely to be an important pathway for BBR’s effects.

Recent studies have focused on the role of the gut-testis axis in spermatogenesis. For example, chronic alcohol-induced dysbiosis of the gut microbiota impairs sperm quality in mice [15]. Dysbiosis of the gut microbiota induced by a high-fat diet causes impaired sperm production and motility [2]. Bacterial overgrowth in the small intestine also impaired spermatogenesis [16]. Alginate oligosaccharides restored spermatogenesis disrupted by busulfan by increasing the abundance of *Bacteroidales* and *Lactobacillaceae* and decreasing the abundance of *Desulfovibrionaceae* [17]. Here, we report that BBR administration impairs spermatogenesis in mice via the gut-testis axis. Mechanistic studies revealed that the decrease in Muribaculaceae abundance in gut microbiota due to BBR administration resulted in reduced ornithine levels, which subsequently decreased the chromatin accessibility of *Ldlr*’s promoter. A decline in LDLR expression hinders LDLR-mediated testosterone synthesis and subsequent spermatogenesis.

## Material and methods

### Animals

ICR male mice of eight weeks old were maintained under a light: dark cycle of 12:12 h at 23±2 °C. The mice had free access to food and water. All animal protocols were approved by the Institutional Animal Care and Use Committee of the Wuhan University. All animal experiments were performed in accordance with the Guidelines for Animal Experiments.

### BBR administration

BBR powder (Macklin) was dissolved in normal saline and shaken. Based on the clinical dose of BBR for diarrhea treatment, three concentration gradients with medium at 120 mg/kg of BBR were tested. Male ICR mice were randomly assigned to four groups (7 mice per group): the control groups (only saline), the lower-dose groups (60 mg/kg BBR), the medium-dose groups (120 mg/kg), and the higher-dose groups (240 mg/kg). After two weeks of intragastric administration, mouse testes, epididymis, feces, and serum samples were collected. Polyethylene glycol (PEG) gavage experiments were performed as described elsewhere [18]. Briefly, 15% PEG 3350 (MiraLax) dissolved in water was administered intragastrically to mice for 2 weeks, and the biological samples were collected as described above.

### Fecal microbiota transplantation and bacteria or metabolite supplementation experiments

Before fecal microbiota transplantation, the mice were pre-treated with a cocktail of broad- spectrum antibiotics (Abx), including 0.2 g/L neomycin sulfate, 0.2 g/L etronidazole, 0.2 g/L ampicile and 0.1 g/L vancomycin, which were added to the drinking water and administered to mice for 2 weeks. Then, 400 mg of fresh stool samples was collected from mice in the control or higher-dose groups and resuspended in 4 mL of saline, vortexed, and filtered.

Finally, 300 μL of the prepared fecal supernatant containing fecal microbiota was intragastrically administered to the mice once every 2 days for 2 weeks.

Bacterial colonization and exogenous metabolite supplementation experiments were performed as described below. The Muribaculaceae representative (*M.intestinale*, DSMZ 28989) was grown on chopped meat at 37 °C anaerobically. A cocktail of *M.intestinale* was resuspended in saline at 3×10^9^/ml. The mice were intragastrically administered 1% ornithine or *M.intestinale* cocktail (200 μL/mouse) once every 2 days for 2 weeks.

### Histological examination

Testes or cauda epididymis were removed from mice, fixed in 4% formalin, embedded in paraffin, and cut into 5μm sections. The sections were then stained with hematoxylin-eosin (H&E) and observed under a light microscope (OLYMPUS IX71).

### Immunofluorescence staining

Testes were removed from mice, fixed in 4% formalin, embedded in paraffin, and cut into 5μm sections. Then sample were permeabilized and blocked. The primary antibodies and secondary antibodies were added to label the target protein, and DAPI was used to seal the slides. Finally, observed under a fluorescence microscope. The antibodies used in this article are listed in Supplementary Table S1.

### Testosterone measurement

Mouse serum was collected, and testosterone levels were detected using an ELISA Kit (ABCLONAL) according to the manufacturer’s instructions. The lowest detection limit for the ELISA kit was 0.19 ng/mL.

### 16S rRNA sequencing of gut microbiota

Fresh feces were collected under sterile conditions, and fecal DNA was extracted using the Magnetic Soil and Stool DNA Kit (TIANGEN), according to the manufacturer’s instructions. Bacterial 16S rRNA gene sequences (V3-V4 region) were amplified using the specific primers 341F:5’-CCTAYGGGRBGCASCAG-3’ and 806R:5’-GGACTACNNGGGTATCTAAT-3’. Sequencing was conducted by the Novogene Technology Company (Beijing, China).

### Extraction of testis RNA and sequencing

Testes were harvested at the indicated time points, as described above. Total RNA was first extracted from the testes, and mRNA was purified using poly T oligo-attached magnetic beads. Fragmentation was carried out using divalent cations and reverse-transcribed into cDNA using random primers. Second-strand cDNA synthesis was subsequently performed using DNA Polymerase I and RNase H. Remaining overhangs were converted into blunt ends, poly(A) was added, and the fragment products were ligated to the Illumina sequencing adaptor. The ligation fragments were size-selected using the AMPure XP system (Beverly, USA), and PCR amplification and sequencing on Illumina platforms with the PE150 strategy were performed by Novogene Bioinformatics Technology (Beijing, China). Differentially expressed genes were screened using the criteria of fold change >1.5 and P-value <0.05.

### Real-time quantitative PCR

Total RNA from the testes or cultured cells was extracted using the Total RNA Isolation Kit (Vazyme) and reverse-transcribed into cDNA using the HiScript II 1st Strand cDNA Synthesis Kit (Vazyme). The primer sets used for quantitative real-time PCR (qRT-PCR) are listed in Supplementary Table S2. qRT-PCR was then performed. Relative gene expression (fold-change) was measured using the 2^−ΔΔCt^ method, in which β-actin was used as the internal control.

### Untargeted metabolomics by liquid chromatography (LC)-MS/MS

Fresh feces (100 mg) were collected and dissolved in methanol. The supernatant obtained after centrifugation was used for LC-MS/MS analysis using a Vanquish UHPLC system (Thermo Fisher Scientific) coupled with an Orbitrap Q Exactive HF-X mass spectrometer (Thermo Fisher Scientific) in both the positive and negative ion modes in Novogene (Beijing, China). Differential metabolites were identified by fold change ≥2 and *P*-value < 0.05.

### Chromatin immunoprecipitation (ChIP) assays

TM3 cells were cross-linked and quenched with glycine. The nuclei were extracted and the DNA was fragmented into 150-300 bp in length by sonication. Lysates were incubated with anti-H3K4me3 antibody or IgG, and the Protein A/G agarose beads (GE Healthcare) were added. Beads were washed once with low-salt buffer, twice with high-salt buffer, twice with LiCl buffer, and twice with TE buffer. The washed beads were eluted, and crosslinking was reversed. After incubation with RNase A and proteinase K, the DNA was purified for qPCR.

### Statistical analysis

For multiple comparisons, one-way ANOVA was performed using SPASS 26.0 software with Duncan’s and LSD methods to identify differences among groups.

## Results

### BBR administration impairs spermatogenesis in mice

Adult mice were intragastrically administered BBR daily for 2 weeks to study the effects of BBR on the male reproductive system. According to the clinical dose for diarrhea treatment, three concentration gradients with medium at 120 mg/kg of BBR were tested. BBR administration significantly caused dose-dependent weight loss in mice (Fig. 1a). However, the testis weights were indistinguishable between control and BBR-treated mice (Fig. 1b).

**Fig. 1.**
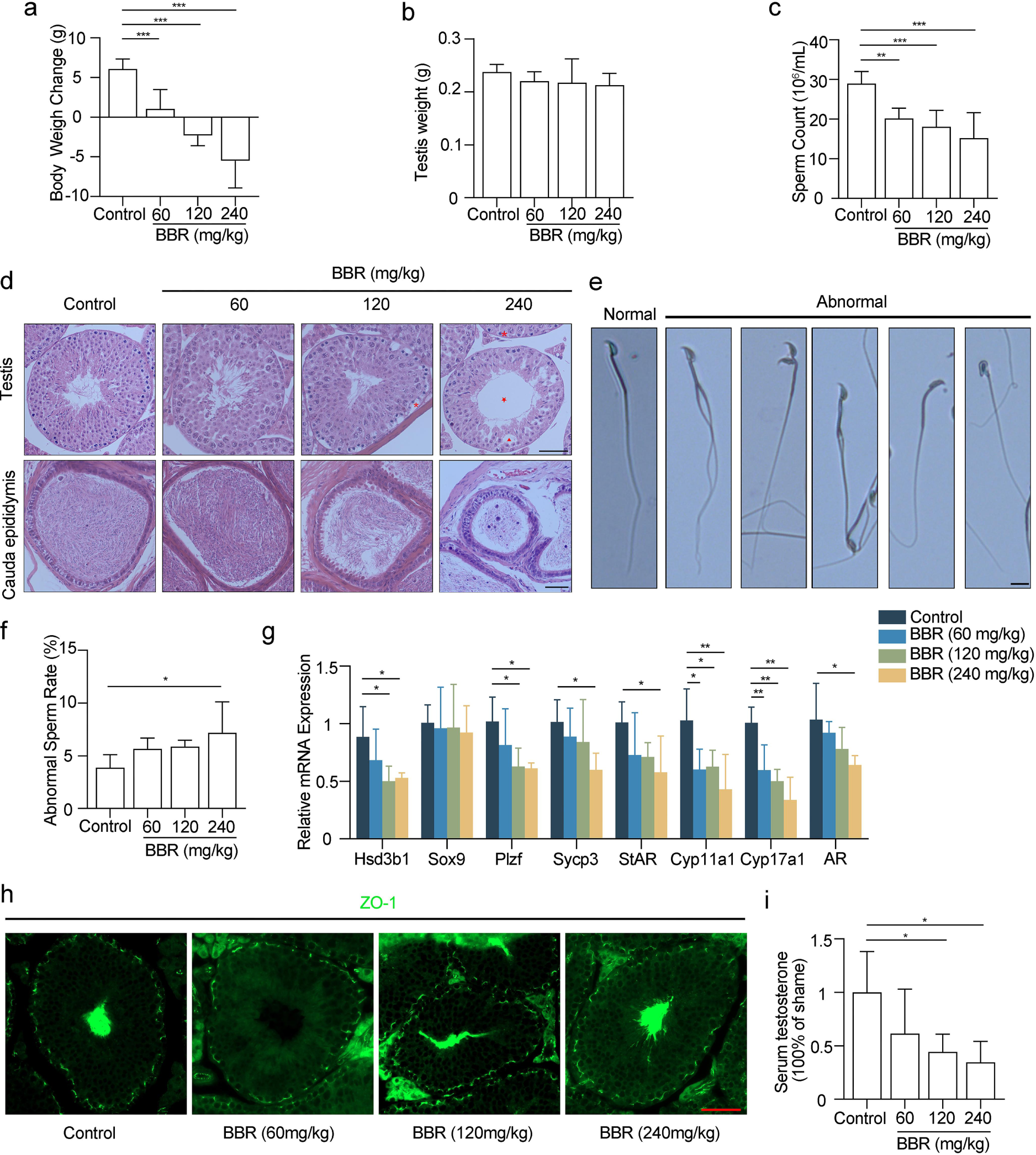
BBR impairs spermatogenesis via attenuating testosterone production. **a** Body weight changes of mice were measured after 14 days of treatment with different BBR dosages. N=7 for each group. **b** Testis weight of mice from control and BBR-treated groups were measured. N=7 for each group. **c** Sperm from cauda epididymis of mice was counted in a haemocytometer under a light microscope. N=7 for each group. **d** Histological analysis of H&E-stained mouse testes and cauda epididymis. Asterisk, vacuolation of spermatogonia. Triangle, disordered arrangement caused by vacuolization of testicular cells. Pentagram, vacuolation of seminiferous tubules. Scale bar, 50μm. **e** Morphological analysis of sperm in mice between control and BBR-treated groups. The spread sperms were stained with Giemsa-staining. Scale bar, 10μm. **f** The rate of sperm deformation in four groups was analyzed. N=4 for each group. **g** Immunofluorescence analysis of BTB integrity. The ZO-1 protein (green) was used to visualized BTB of the testes. Scale bar, 50μm. **h** The mRNA expression levels of marker genes of testicular cells, genes related to testosterone synthesis and AR were measured. N=4 for each group. **i** The testosterone of mouse serum was measured by enzyme linked immunosorbent assay (ELISA). N=4 for each group. Data are expressed as mean±SEM. Differences without statistical significance were not labled, *p<0.05, **p<0.01, ***p <0.001.

We found that sperm concentrations in the epididymis decreased significantly with increasing BBR dosage (*P<0.01*) (Fig. 1c). Histological examination by hematoxylin-eosin (H&E) staining of the testes and epididymis of mice showed that mice in the control groups had normal testicular architecture and germinal cell arrangement, and mice in the lower dose groups had no distinguishable morphological differences in the testicular sections when compared with the controls (Fig. 1d). However, mice in the medium-dose and higher-dose groups showed disordered arrangement and vacuolation in the seminiferous epithelium of the testis, which was even worse in the higher-dose groups (Fig. 1d). Apoptotic sperm plaques appeared in the epididymis of the medium-dose and higher-dose groups and were worse in the higher-dose mice (Fig. 1d). Mice exposed to a higher dose of BBR had a higher sperm deformity rate in the epididymis (*P<0.01*) (Fig. 1e-f). As shown in Fig. 1e, the sperm deformity types included double tails, double heads, double necks, banana-shaped heads, and irregular heads. Except for Sertoli cells gene *Sox9*, the mRNA levels of Leydig cells gene *Hsd3b1*, spermatogonia gene *Plzf* and spermatocytes gene *Sycp3* were down-regulated with increase of BBR dose (Fig. 1g). The blood-testis barrier (BTB), as a gatekeeper, protects the developing germ cells of males [19]. Immunofluorescence showed that the tight junction protein constructed by ZO-1 were not disrupted in any of the groups (Fig. 1h). The results strongly declared that BBR disrupted spermatogenesis by affecting Leydig cells. Besides, BBR treatment caused a dose-dependent decrease in serum testosterone levels in mice (Fig. 1i). Meanwhile, the mRNA levels of *StAR*, *Cyp11a1*, *Cyp17a1*, which are related to testosterone synthesis, decreased with increasing BBR dosage (Fig. 1g). In response, androgen receptor (AR) also decreased (Fig. 1g). Taken together, these results reveal that BBR administration disrupts spermatogenesis by influencing testosterone synthesis of Leydig cells in the testes.

### BBR administration alters gut microbiota composition

The variation of gut microbiota in mice was focused on because of the poor absorption and low bioavailability of BBR [20]. We sequenced the bacterial 16S rRNA gene in the feces. The sparse curves of bacterial communities reached a saturation plateau, indicating a sufficient sequencing depth for detecting the majority of gut bacteria (Supplementary Fig. S1). The total bacterial load or category decreased slightly with increasing BBR dosage (Fig. 2a). In addition, the gut microbiota richness represented by the Chao1 index and diversity represented by Shannon and Simpson indices were destroyed with increasing BBR dosage according to microbial alpha analysis (Fig. 2b-d). Microbial beta diversity analysis based on Principal Component Analysis (PCA) showed obvious differences in the gut microbiota composition between control and BBR-treated mice (Fig. 2e). The ratio of Firmicutes to Bacteroidota was not significantly different (Fig. 2f). We then analyzed the changes in the abundance of the dominant bacteria. At the phylum level, medium-dose and higher-dose of BBR treatment significantly depleted Bacteroidota in the gut of mice, and increased Proteobacteria abundance was detected in the higher-dose groups (Fig. 2g). At the order level, higher-dose of BBR eliminated Bacteroidales but enriched Lachnospirales (Fig. 2h). At the genus level, a depletion of *Muribaculaceae* and an enrichment of *Bacteroides* were observed in the medium-dose and higher-dose groups, and the higher-dose of BBR also apparently enriched *Robinsoniella* (Fig. 2i). In addition, BBR significantly altered the abundance of the bacterial families (Fig. 2j). The predominant bacteria in the control groups were Muribaculaceae, Lactobacillaceae, and Bacteroidaceae. In contrast, the predominant bacteria in the higher-dose groups were Bacteroidaceae, Lachnospiraceae, and Enterobacteriaceae. Remarkably, the relative abundance of Muribaculaceae decreased from 55.77% to 5.63% in the higher-dose groups. Line discriminant effect size (LEfSe) analysis identified these bacteria with differential abundance (Fig. 2k). In summary, these data revealed that BBR altered the composition and relative abundance of gut microbiota, which prompted us to ask whether dysbiosis of the gut microbiota disrupted spermatogenesis in mice.

**Fig. 2.**
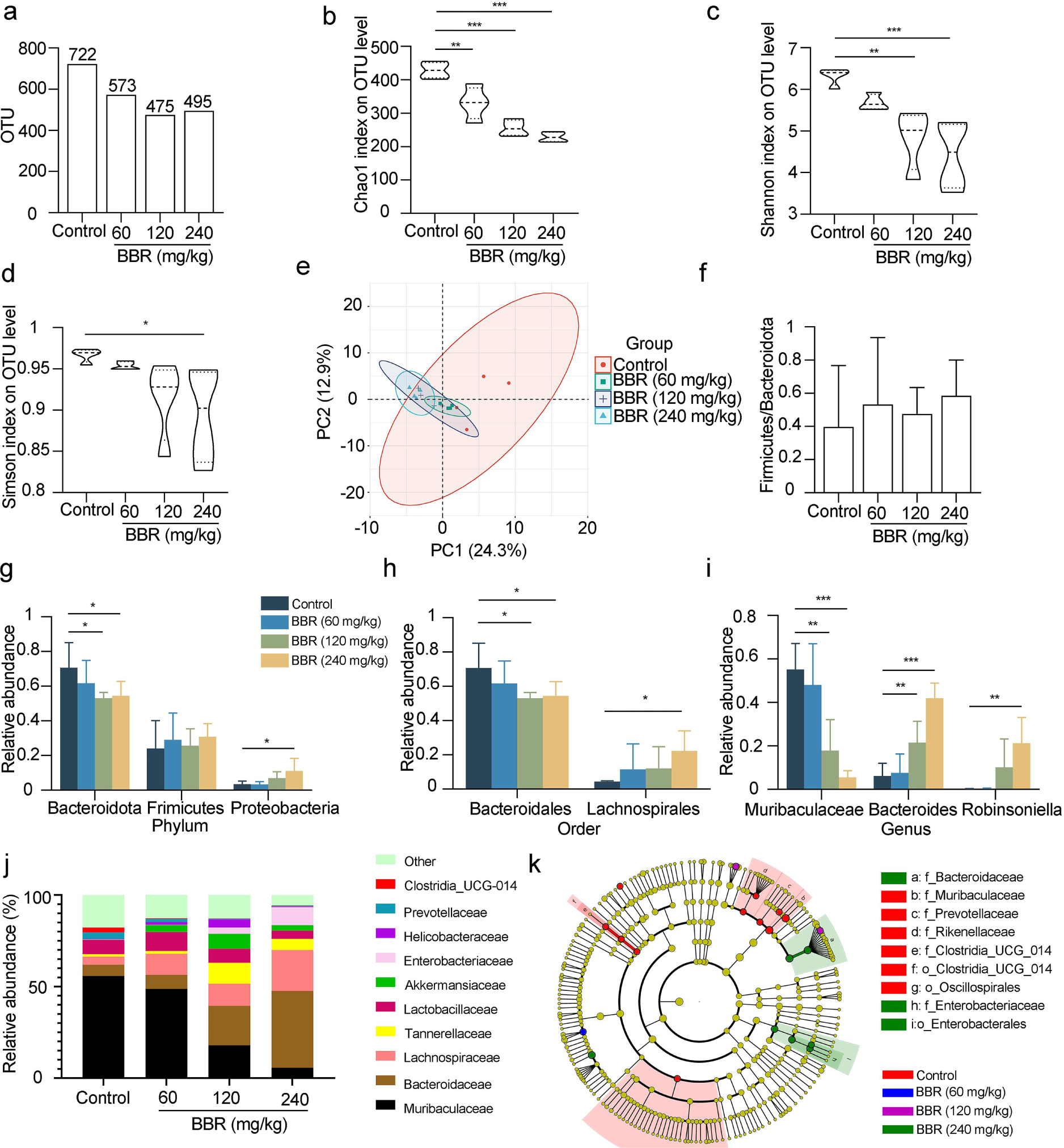
BBR induced gut microbiota dysbiosis in mice. **a** The total fecal bacterial load and category were analyzed in mice with different BBR dosages. N=6 for each group. **b-d** The Chao1 index (**b**), Shannon index (**c**) and Simpson index (**d**) were analyzed. N=6 for each group. **e** Principal component analysis (PCA) plot was generated based on the similarity for the samples in the untreated and BBR-treated groups. N=6 for each group. **f** The ratios of Firmicutes/Bacteroidota were analyzed at the phylum. N=6 for each group. **g-i** The relative abundances of dominant bacteria at phylum (**g**), order (**h**) and genus (**i**) levels were analyzed. N=6 for each group. **j** The relative taxonomic abundances at the family level of gut microbiota were showed as a histogram. N=6 for each group. **k** Taxonomic cladogram from linear discriminant analysis effect size (LEfSe) was showed. Dot size is proportional to the abundance of the taxon. N=6 for each group. Data are expressed as mean±SEM. Differences without statistical significance were not labled, *p<0.05, **p<0.01, ***p <0.001.

### Muribaculaceae is responsible for BBR’s effects on spermatogenesis in mice

To verify the effect of gut microbiota on spermatogenesis, mice pretreated for gut microbiota reconstitution with an antibiotic cocktail were used for faecal microbiota transplantation (FMT) with BBR-treated or control donors (Fig. 3a). Body weight and epididymal sperm concentration were significantly reduced in BBR-FMT mice, strongly indicating that gut microbiota may regulate spermatogenesis in mice (Fig. 3b-c). The 16S rRNA sequencing of gut microbiota showed a drastic decrease in the Muribaculaceae abundance in BBR-treated mice, followed by the correlation of Muribaculaceae and sperm concentration in mice (Fig. 3d). As expected, the relative abundance of Muribaculaceae was highly correlated with the sperm concentration (r=0.7352, *P*=0.0012). Polyethylene glycol (PEG) can alter intestinal osmolality and deplete Muribaculaceae abundance [18]. Therefore, PEG administration experiments were performed to verify the role of Muribaculaceae in spermatogenesis (Fig. 3e). The total bacterial abundance or taxa in the feces decreased significantly after PEG treatment (Supplementary Fig. S2a-d). Microbial beta diversity analysis based on PCA showed obvious differences in the gut microbiota composition between the control and PEG treatment groups (Fig. 3f). Although the richness and diversity of microbes were reduced in the intestines of PEG-treated mice, the ratio of Firmicutes to Bacteroidota was not significantly different between control and PEG-treated mice (Supplementary Fig. S2e). As expected, the dominant microbiota of the intestine in mice treated with PEG was converted from Muribaculaceae to Bacteroidaceae at the family level (Fig. 3g), and the relative abundance of Muribaculaceae decreased from 53.42% to 1.79% (Fig. 3h, Supplementary Fig. S2f). Histological examination showed that mice in the PEG-treated groups had a lower epididymal sperm density, dropping by 46.73% (Fig. 3i-j). To determine the upstream factor causing sperm reduction, we collected serum from PEG-treated and control mice and performed an ELISA against testosterone; Similar to BBR-treated mice, the serum testosterone levels in the PEG-treated groups decreased by 42.27% when compared with the control groups (Fig. 3k). In addition, the mRNA levels of *StAR*, *Cyp11a1* and *Cyp17a1* in the testes were significantly decreased in the PEG-treated mice (Fig. 3l). Thus, the decrease in Muribaculaceae was responsible for attenuated sperm production, and the effect worked by regulating testosterone levels.

**Fig. 3.**
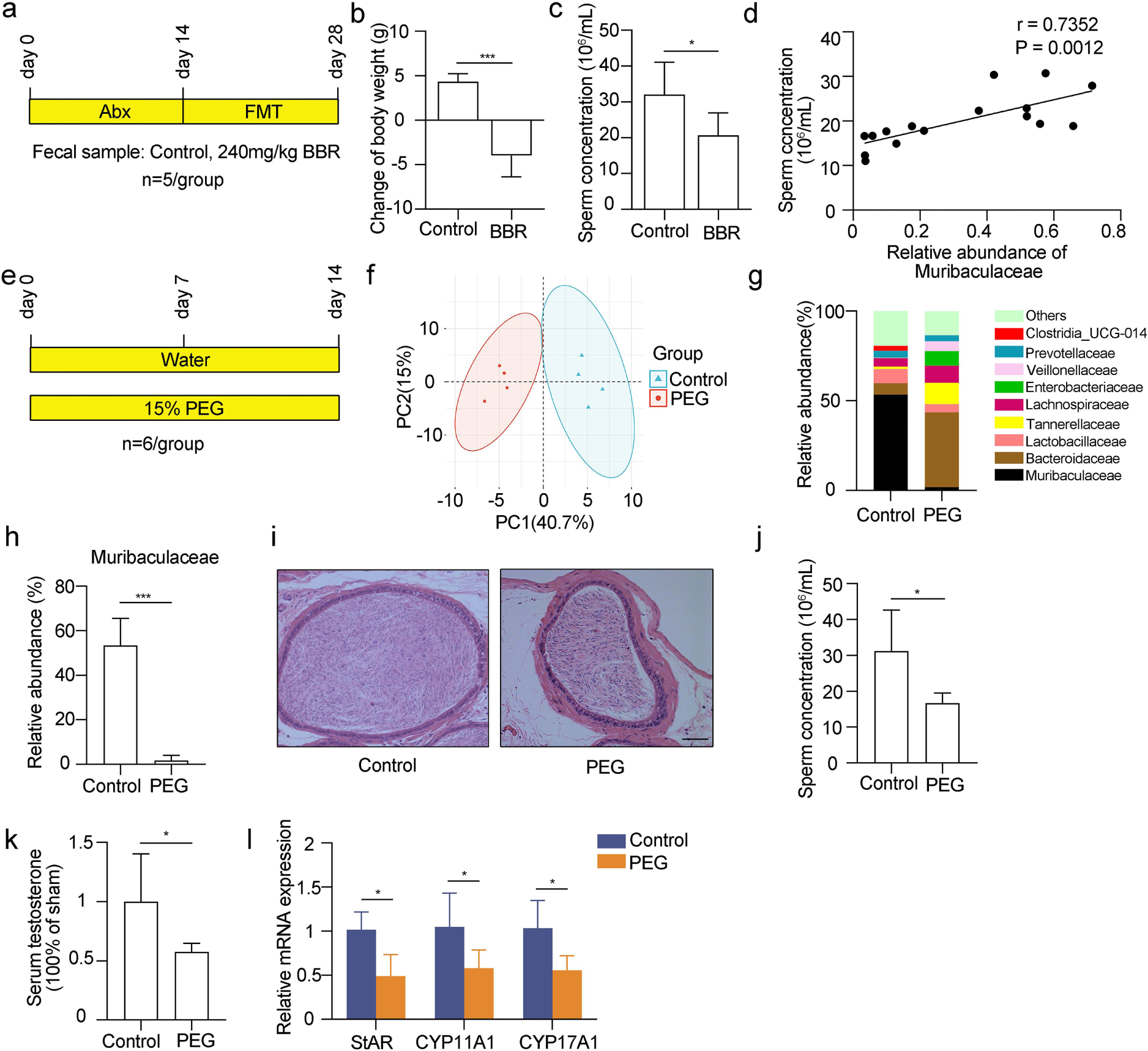
T**h**e **adverse effects of BBR on spermatogenesis could be transferred by FMT. a** Schematic diagram of faecal microbiota transplantation experiment. Abx, a cocktail of broad-spectrum antibiotic. N=5 for each group. **b** Statistical analysis of changes in body weight after FMT. **c** Sperm concentrations from cauda epididymis of mice were analyzed in a haemocytometer under a light microscope. **d** The correlation between Muribaculaceae abundance and sperm concentration was analyzed in control, lower-dose, medium-dose and higher-dose groups. **e** Schematic diagram of PEG treatment experiment. N=6 for each group. **f** PCA plot was generated based on the similarity for faecal samples in untreated and PEG-treated groups. **g** The relative taxonomic abundances at the family level of gut microbiota were showed as a histogram. **h** The relative abundance of *Muribaculaceae* in intreated and PEG-treated groups was analyzed. N=4 for each group. **i** Histological analysis of mouse cauda epididymis. The sections were stained with H&E. Scale bar, 50μm. **j** Sperms from cauda epididymis of mice were counted in a haemocytometer under a light microscope, N=6 for each group. **k** The testosterone of mouse serum in untreatedand PEG-treated was measured by enzyme linked immunosorbent assay (ELISA). N=5 for each group. **l** The mRNA expression levels of genes related to testosterone synthesis were measured by RT-qPCR. N=4 for each group. Data are expressed as mean±SEM. Differences without statistical significance were not labled, *p<0.05, **p<0.01, ***p <0.001.

### BBR alters gut microbiota metabolism

The gut microbiota regulates the biological processes of the host through its metabolites, prompting us to ask whether Muribaculaceae disorder causes a decrease in testosterone levels through its metabolites. Thus, we uncovered metabolic features of the gut microbiota of mice in the BBR-treated and control groups by liquid chromatography-mass spectrometry (LC-MS/MS). PCA score plots showed that the higher-dose BBR-treated groups were clearly separated from the control groups (Fig. 4a). We then compared the faecal metabolomic profiles of both groups and found 394 downregulated metabolites and 284 upregulated metabolites in the BBR-treated groups when compared with the control groups (Fig. 4b).

**Fig. 4.**
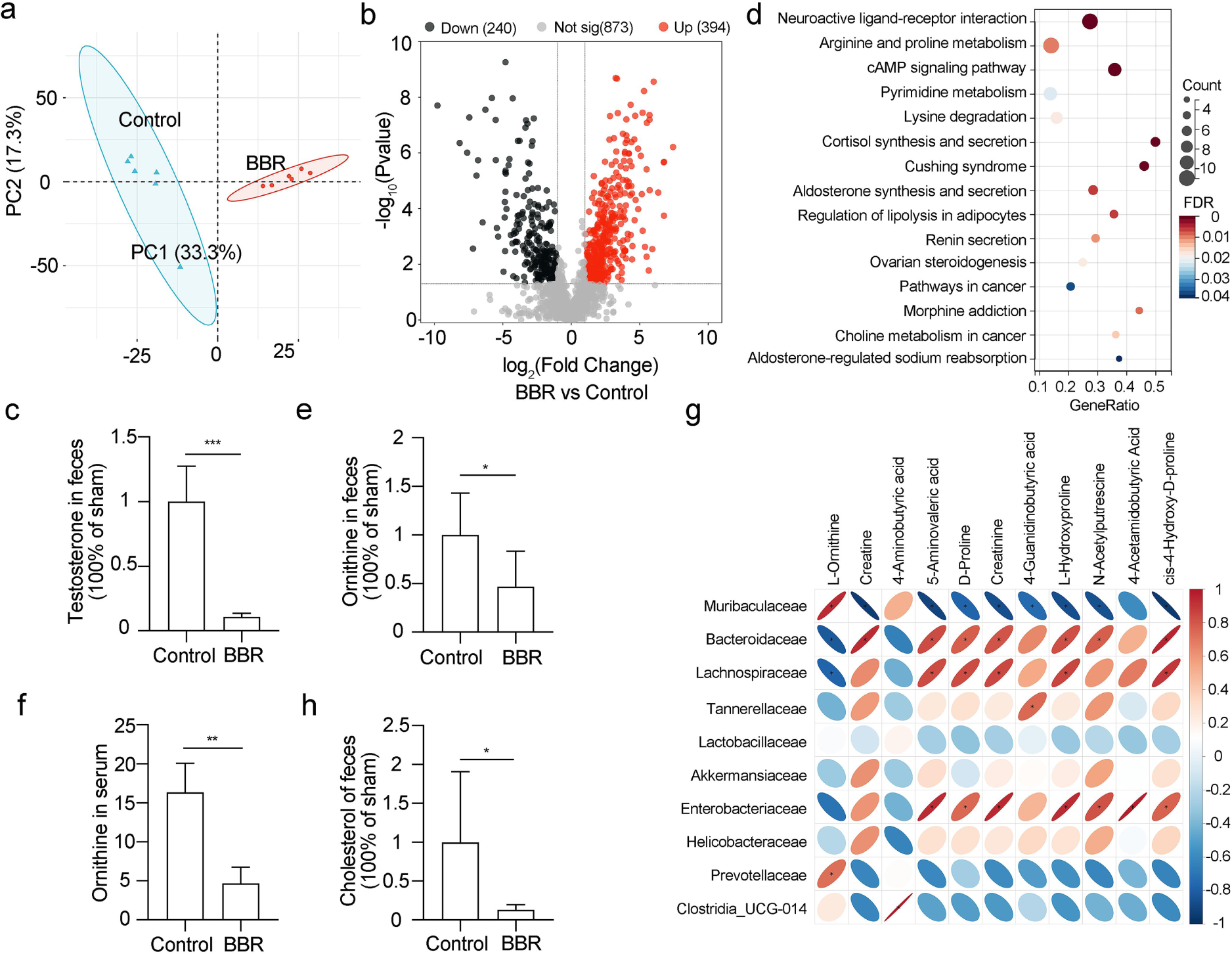
BBR affected ornithine metabolism of mice gut microbiota. **a** PCA plot was generated based on the similarity for the samples in the untreated and BBR- treated (higher-dose) groups. N=6 for each group. **b** The differential metabolites of gut microbiota were showed with volcano plot. P-value was <0.05. **c** The testosterone levels of faecal samples were analyzed according to LC-MS/MS-based metabolomic results. **d** Significant Kyoto Encyclopedia of Genes and Genomes (KEGG) pathways for the fecal microbiome of untreated and BBR-treated groups were identified. **e** The ornithine levels of feces were analyzed. N=4 for each group. **f** The ornithine levels of mice serum were measured by high performance liquid chromatography (HPLC). N=3 for each group. **g** The close correlations between bacteria abundance at family levels and the differential metabolites of arginine and proline metabolism pathway were analyzed in untreated and BBR-treated groups.

Similar to the testosterone level in the serum, the level of fecal testosterone was also decreased by 89.17% when compared with the controls, which illustrated that testosterone synthesis from Leydig cells was disrupted (Fig. 4c). Differential metabolites were enriched to identify the biological pathways involved, including neuroactive ligand-receptor interaction, arginine and proline metabolism, cAMP signaling pathway, pyrimidine metabolism, and lysine degradation (Fig. 4d). Interestingly, Muribaculaceae is related to the ornithine biosynthesis pathway by annotation of bacterial genomes [21], and ornithine is an indispensable metabolite in the arginine and proline metabolism pathways, which prompted us to explore the correlation between Muribaculaceae and ornithine. To verify this hypothesis, we performed a correlation analysis between the top 10 microbes at the family level and differential metabolites of arginine and proline metabolism pathways (Fig. 4e). The results showed that Muribaculaceae was highly correlated with metabolites in the arginine and proline metabolism pathways, including L-ornithine (r=0.9313), creatine (r=-0.9344), 4- aminobutyric acid (r=0.5035), 5-aminovaleric acid (r=-0.9013), D-proline (r=-0.7701), creatinine (r=-0.8688), 4-guanidinobutyric acid (r=-0.7406), L-hydroxyproline (r=-0.8694), N-acetylputrescine (r=-0.8534), 4-acetamidobutyric acid (r=-0.5857), and cis-4-hydroxy-D-proline (-0.9467). We observed that Muribaculaceae and L-ornithine were highly positively correlated, and ornithine levels in the feces and serum of mice in the BBR-treated groups decreased drastically (Fig. 4f-g). Taken together, we conclude that Muribaculaceae may regulate ornithine synthesis.

### *M.intestinale*-produced ornithine is necessary for spermatogenesis in mice

To verify our hypothesis, we cultured Muribaculaceae *in vitro* and investigated its function. There are ten species in the Muribaculaceae family, and *M.intestinale*, which are representative of four isolates [22], were cultured anaerobically in *vitro* (Supplementary Fig. S3). Annotation of genomic function indicated that acetylglutamate kinase (ArgB) and N- acetylglutamylphosphate reductase (ArgC), two important enzymes involved in the conversion of ornithine to glutamate, were specifically expressed in *M.intestinal* genome (Fig. 5a). Therefore, we detected glutamate and ornithine levels in the medium after *M.intestinale* culture for 24 h. Compared with the pre-cultured medium without *M.intestinale*, glutamate levels decreased and ornithine levels increased in the medium after *M.intestinale* culture (Fig. 5b-c). These results strongly confirmed that *M.intestinale* can utilize glutamate as a substrate to produce ornithine.

**Fig. 5.**
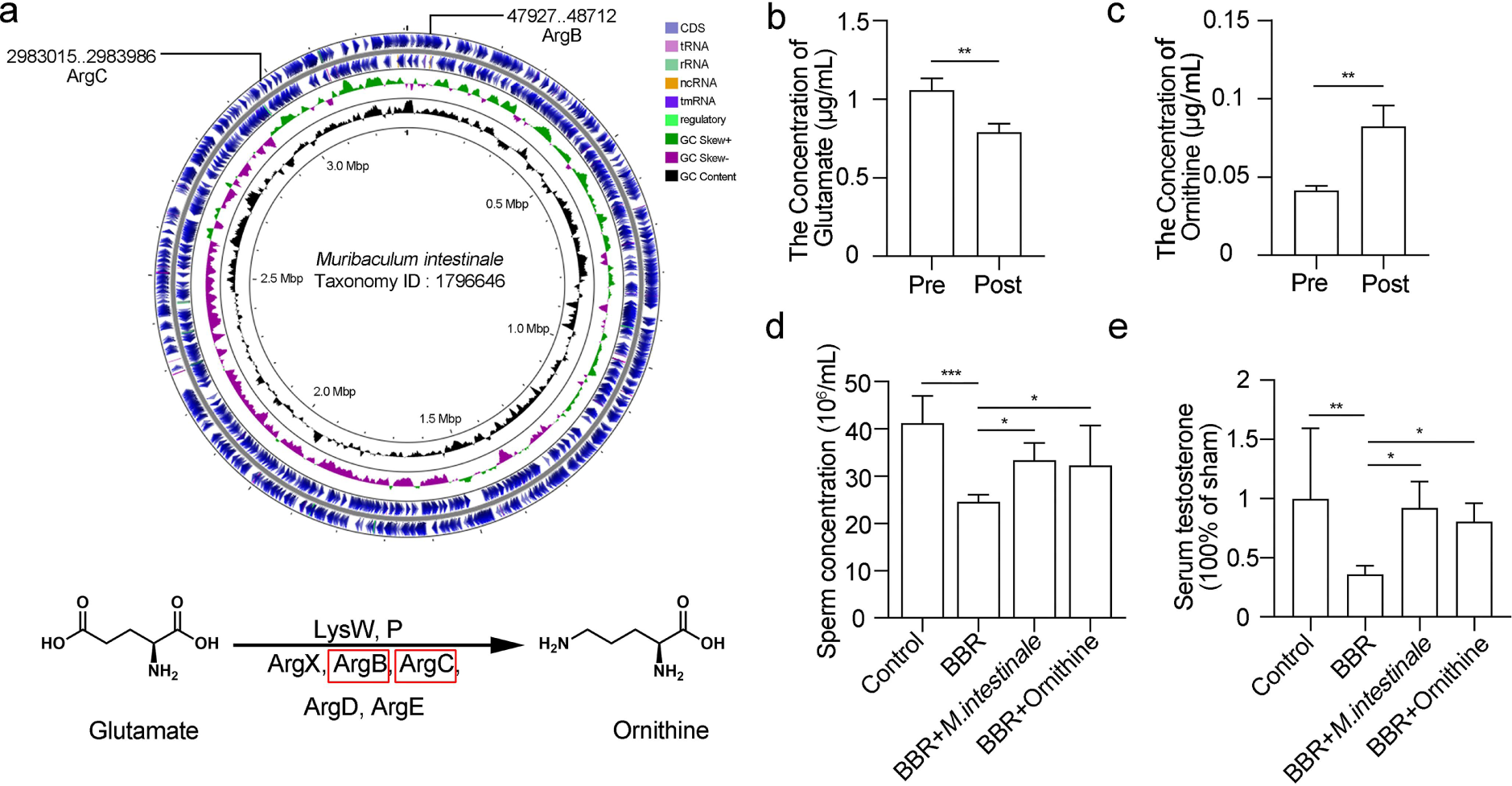
Oral administration of *M.intestinale* or ornithine rescued BBR-induced spermatogenic defects. **a** (Top) Structure and distribution of ArgB, ArgC for ornithine synthesis on the chromosome of *M. intestinale* were showed. The proposed gene products are identified in straight lines. (Down) The pathway of the conversion of glutamate to ornithine was demonstrated. **b-c** The glutamate levels (**b**) and ornithine levels (**c**) in the medium with or without *M. intestinale* were measured by HPLC. *M.intestinal* was cultured in vitro for 24h. Pre, pre-cultured. Post, post-cultured. **d** The sperm concentrations of untreated, BBR-treated, BBR+*M.intestinale*- treated, BBR+ornithine-treated mice were counted in a haemocytometer under a light microscope, N=5 for each group. **e** The testosterone level of mouse serum was measured by ELISA. N=5 for each group. Data are expressed as mean±SEM. Differences without statistical siganificance were not labled, *p<0.05, **p<0.01, ***p <0.001.

The decrease in Muribaculaceae abundance was irreversible, which prompted us to ask whether spermatogenesis disruption resulting from Muribaculaceae depletion could be reversible. The *M.intestinale* culture medium or ornithine was intragastrically administered to mice for 14 days when BBR was simultaneously fed to mice, and the sperm concentration and testosterone levels were measured. As expected, administration of either *M.intestinale* or ornithine reversed the decline in sperm concentration in mice treated with BBR by 88.11% and 85.19%, respectively (Fig. 5d). Meanwhile, testosterone levels also recovered to a certain extent, with mice treated with BBR and *M.intestinale* recovering to 92.23%, and mice treated with BBR and ornithine recovering to 80.69% of that of untreated mice (Fig. 5e). These data suggest that *M.intestinale* and ornithine administration could prevent the disruption caused by BBR during spermatogenesis.

### BBR influences steroid metabolic process

To further delineate the mechanism by which the metabolite ornithine of *M.intestinale* regulates spermatogenesis, we collected whole testis samples for RNA sequencing. A total of 672 differentially expressed genes were identified, with an expression change greater than 1.5-fold between the higher-dose BBR-treated groups and control groups. Among them, 314 genes were upregulated and 358 genes were downregulated after BBR treatment (Fig. 6a).

**Fig. 6.**
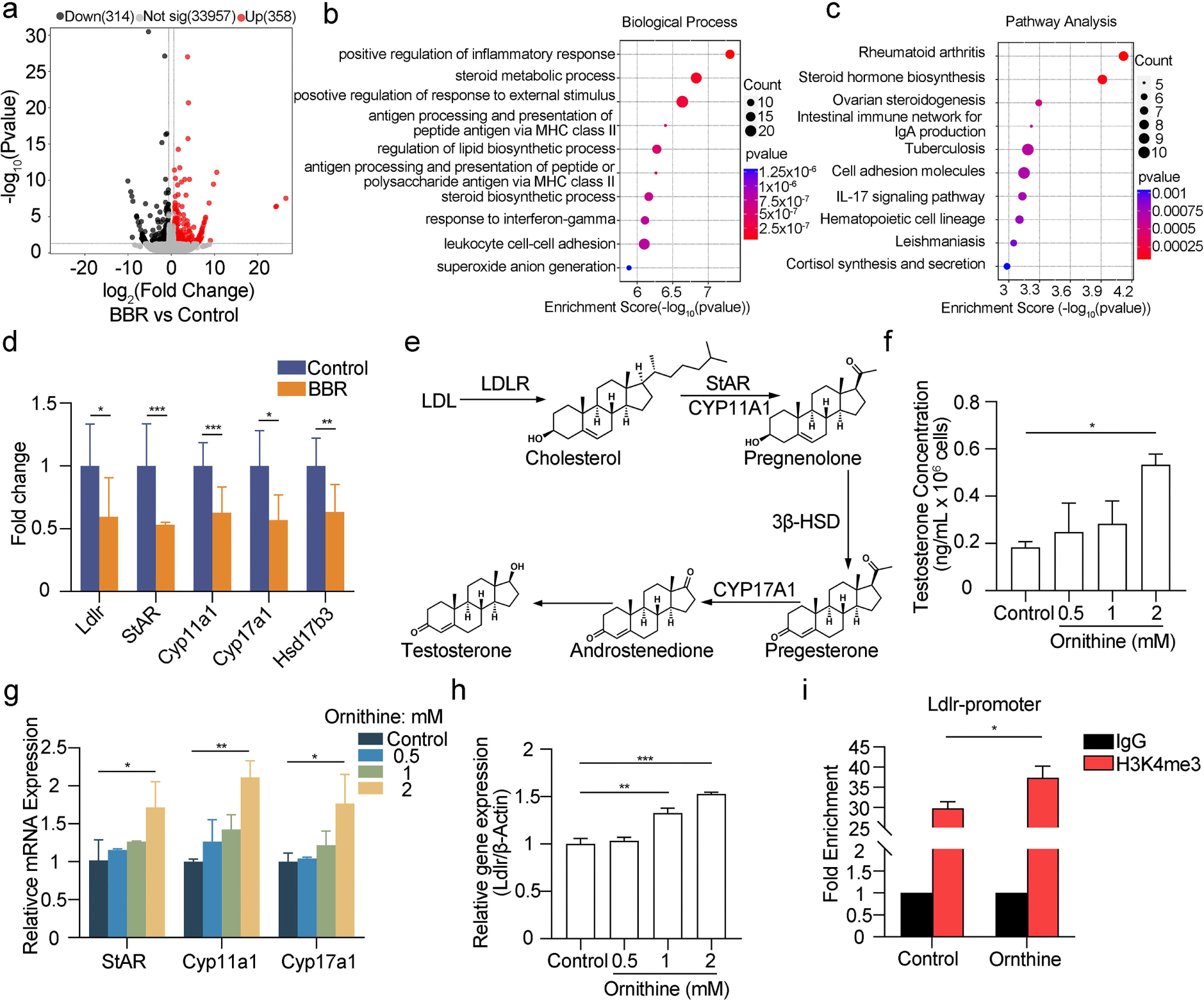
BBR affected the steroid metabolic process related to *Ldlr* in mice testis, whose promoter accessibility was promoted by ornithine. **a** The differentially expressed genes between control and BBR-treated (higher-dose) mice were analyzed. P-value<0.05. **b** The biological process involved in differentially expressed genes were analyzed through gene ontology (GO). **c** Significant Kyoto Encyclopedia of Genes and Genomes (KEGG) pathways for the differentially expressed genes were identified. **d** The mRNA levels of *Ldlr*-associated genes related to testosterone synthesis were analyzed. N=2 for each group. **e** LDLR-induced testosterone synthesis pathway was showed. **f** The cholesterol levels of feces were analyzed. N=6 for each group. Data are expressed as mean±SEM. Differences without statistical siganificance were not labled, *p<0.05, **p<0.01, ***p <0.001. **g** The testosterone levels of medium were measured after TM3 were treated with different concentrations of ornithine (0. 0.5, 1 and 2mM). N=2 for each group. **h** The mRNA expression levels of genes related to testosterone synthesis were measured by RT- qPCR. N=2 for each group. **i** *Ldlr* mRNA levels were measured by RT-qPCR. N=2 for each group. **j** ChIP-qPCR assay for the direct binding of H3K4me3 to *Ldlr* promoter was performed. IgG was used as the negative control. N=3 for each group. Data are presented as mean±SEM. Differences without statistical significance were not labled, *p<0.05, **p<0.01, ***p <0.001.

Gene ontology (GO) analysis of these differentially expressed genes revealed a series of biological processes, such as the positive regulation of inflammatory response, positive regulation of response to external stimulus, antigen processing and presentation of peptide antigen via MHC class II, and regulation of lipid biosynthetic process, particularly the testosterone synthesis-related steroid metabolic process (Fig. 6b). Molecular function analysis also pointed to the steroid-binding function, which is related to testosterone synthesis (Supplementary Fig. S4). Kyoto Encyclopedia of Genes and Genomes (KEGG) pathway analysis showed a series of pathways were involved, such as rheumatoid arthritis, steroid hormone biosynthesis, ovarian steroidogenesis, intestinal immune network for IgA production and tuberculosis (Fig. 6c). Among these, steroid hormone biosynthesis is critical for testosterone synthesis. In particular, a series of testosterone synthesis-related genes were significantly downregulated, including *Ldlr*, *StAR*, *Cyp11a1*, *Cyp17a1* and *Hsd17b3* (Fig. 6d). Testosterone synthesis in Leydig cells was mainly regulated by the upstream gene *Ldlr* (Fig. 6e). LDLR transports low-density lipoprotein into Leydig cells to participate in cholesterol synthesis and regulate cholesterol homeostasis [23]. Cholesterol, as a critical substrate, can be used to synthesize testosterone in Leydig cells [24]. Importantly, the cholesterol levels in the feces of BBR-treated mice decreased drastically compared to control mice (Fig. 6f). Based on the above, we speculated that *Ldlr* is a potential target of the metabolite ornithine for lowering cholesterol and testosterone synthesis.

### Ornithine participates in steroid metabolism via regulating Ldlr transcription

Next, we explored how ornithine regulates *Ldlr* expression in TM3 cells, a mouse Leydig cell line. We found that *Ldlr* gene significantly increased with increasing ornithine dosage (Fig. 6g). Meanwhile, the levels of testosterone and genes related to testosterone synthesis increased as ornithine concentration increased (Fig. 6h-i). These data illustrate that ornithine facilitates the expression of *Ldlr* to increase testosterone levels. Histone modifications regulate gene expression by altering chromatin accessibility; histone H3 lysine 4 trimethylation (H3K4me3) and histone H3 lysine 27 trimethylation (H3K27me3) are associated with active and repressive transcription, respectively [25]. Therefore, we performed ChIP-qPCR and analyzed H3K4me3 occupancy of *Ldlr* promoter. Our quantitative analysis revealed a 29.77-fold enrichment of H3K4me3 compared with IgG at *Ldlr* promoter (Fig. 6j). We also detected a 1.25-fold increase in H3K4me3 occupancy in the target region in ornithine-treated groups when compared with the control groups (Fig. 6j), suggesting that ornithine increased the promoter accessibility of *Ldlr* and promoted LDLR expression.

## Discussion

In the present study, we revealed that the dysbiosis of gut microbiota induced by BBR is one of the primary causes of impaired spermatogenesis in the mouse host. As shown in the schematic diagram in Fig. 7, BBR altered the gut microbiota composition, and the abundance of Muribaculaceae drastically decreased, which resulted in the arrest of glutamate conversion due to the loss of ArgB and ArgC expressed by Muribaculaceae in mice. The decrease in ornithine levels led to a decrease in chromatin accessibility, where *Ldlr* promoter was located. Therefore, low-density lipoprotein outside the Leydig cells could not be transported by LDLR into the cytoplasm for intracellular synthesis of cholesterol as a substrate for testosterone, resulting in impaired spermatogenesis due to the decreased testosterone.

**Fig. 7.**
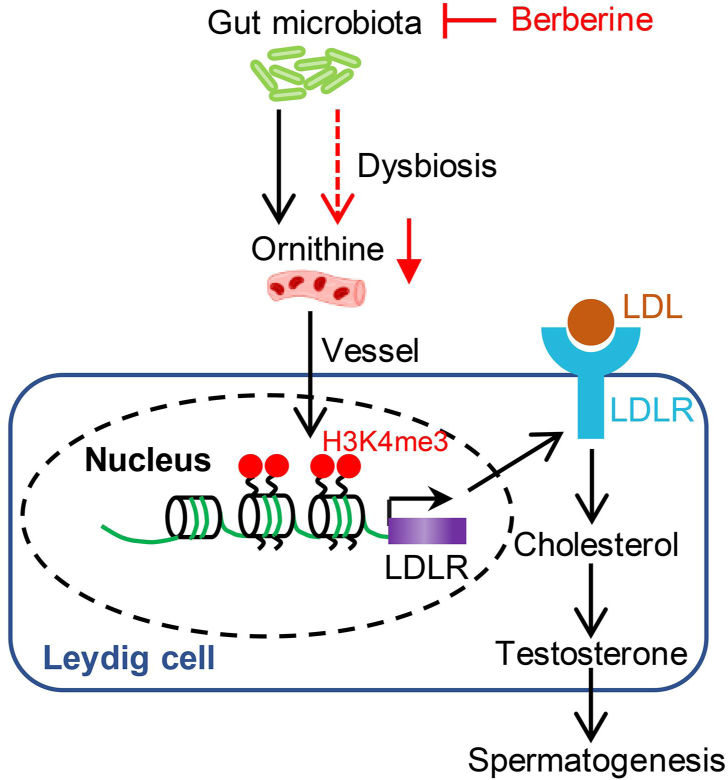
Illustration of the findings that BBR disrupted the spermatogenesis by inducing gut microbiota dysbiosis in mice. LDL, low density lipoprotein. LDLR, low density lipoprotein receptor. H3K4me3, histone H3 lysine 4 trimethylation.

Considering the results, adult couples should avoid taking BBR or drugs containing BBR as much as possible while trying to conceive, and the fertility intention of male patients should be considered when BBR is used to treat diseases.

We noticed that the testicular structure was destructed, and disordered arrangement and vacuolation in the seminiferous epithelium occurred in testes of BBR-treated mice. The results of RNA sequencing showed that inflammatory and immune pathways were affected, which means that inflammation occurred in the testes of BBR-treated mice (Fig. 6b-c). It has been reported that the pro-inflammatory macrophage cells were presented in testes, and testicular inflammation affected sperm quality [26, 27]. We noticed that these vacuolation in the seminiferous epithelium were closer to the seminiferous tubule wall, which was due to inflammation-mediated apoptosis of spermatogonia.

Although BBR has become a drug in treating diarrhea, diabetes, hyperlipidemia, and cancer, it become a question that how it achieves its effects *in vivo* due to the poor bioavailability of BBR for oral administration. It is reported that the intestinal absorption of BBR is less than 1%, and the absorbed berberine could be excreted back to the intestinal lumen with the action of P-glycoprotein [28]. Thus, gut microbiota is the primary pathway for BBR to function. In this study, we found that the abundance of Muribaculaceae and the decline in sperm concentration in mice were closely related. Regarding the drastic decrease in the abundance of Muribaculaceae after BBR treatment in mice, the following explanation seems reasonable: first, BBR can attenuate bactericidal activity and inhibit the growth of protozoa [29]. BBR has been shown to be bactericidal against *V.cholera* and protozoacidal against *Giardia lamblia* [30, 31]. Our results showed a 50.14% decrease in the relative abundance of Muribaculaceae after BBR treatment, indicating that BBR may also exert bactericidal activity against Muribaculaceae family. Second, Muribaculaceae family is highly sensitive to osmolality, and an increase in osmolality leads to a decrease in Muribaculaceae abundance [18]. Polyethylene glycol (PEG), as a compound that cannot be absorbed by the intestinal epithelium, is used to induce changes in osmolality [18], and this is why PEG specifically lowers Muribaculaceae. Therefore, BBR administration may change intestinal osmolality, causing Muribaculaceae to disappear without adapting to the intestinal environment. Notably, the decrease in Muribaculaceae abundance was irreversible.

Although Muribaculaceae is abundant, up to approximately 50% at the family level in the gut of mice [18], there has been little research on Muribaculaceae. Muribaculaceae was speculated to be involved in ornithine biosynthesis via genome annotation, but this hypothesis has not been validated [21]. Correlation analysis revealed a strong correlation between sperm concentration and the abundance of Muribaculaceae. We revealed that Muribaculaceae could synthesize ornithine from glutamate anaerobically *in vitro* and found the biological significance of Muribaculaceae for the first time. These results are significant because when the abundance of Muribaculaceae is downregulated, diseases caused by ornithine deficiency could be considered for detection and diagnosis, or diseases with decreased ornithine levels could be treated by Muribaculaceae supplementation via fecal transplantation.

Notably, the metabolites ornithine is recognized as a bridge to connect the gut microbiota and spermatogenesis by blood circulation, and the decrease in metabolites ornithine level of gut microbiota disrupt spermatogenesis in mouse in our study. For one thing, the causal relationships between the gut microbiome and blood metabolites have been demonstrated.

The metabolites of gut microbiota affect its host, and nearly half of blood metabolites are associated with gut microbial species and their metabolites [32]. The gut microbiota could mediate the transportation of its metabolites through gut barrier into the bloodstream [32]. It has been reported that the host harboring microbes that convert cholesterol into sterol coprostanol has lower fecal cholesterol levels and lower serum total cholesterol [33].

Interestingly, we also found that the fecal ornithine levels decreased significantly in BBR- treated mouse, which illustrated that the ornithine levels in the blood and testis of BRR- treated mice are also decreased to a certain degree (Fig. 4f). For another, polyamines have been reported be involved in cell differentiation and proliferation, and ornithine decarboxylase (ODC1), as the rate-limiting enzyme, metabolized ornithine to polyamines [34]. We found that the mRNA level of *Odc1* decreased significantly in testis, which meaned that the levels of ornithine, the upstream material for polyamines, decreased in testis (Supplementary Fig. S5a).

The biosynthesis of testosterone from cholesterol in Leydig cells plays an essential role in the maintenance and regulation of male fertility. Cholesterol is metabolized to pregenenolone via the inner mitochondrial membrane of CYP11A1, which is subsequently converted to testosterone by mitochondria and smooth endoplasmic reticulum enzymes [35]. Cholesterol is derived from four sources in Leydig cells: de novo synthesis in the endoplasmic reticulum, utilization of cholesteryl esters in lipid droplets by cholesteryl ester hydrolase, uptake of plasma lipoprotein-derived cholesteryl esters by LDLR or SR-BI, and plasma membrane-associated free cholesterol [36]. LDLR-transported low-density lipoprotein is the main source of intracellular cholesterol synthesis [35]. We found that a decline in LDLR expression greatly affects cholesterol synthesis. It has been reported that cold shock domain-containing protein E1 (CSDE1) facilitates degradation of *Ldlr* mRNA [37]. However, we found no difference in the mRNA level of CSDE1 after BBR treatment, indicating that the decline in LDLR expression was not mediated by CSDE1 (Supplementary Fig. S5b). Therefore, we speculate that ornithine regulates the *Ldlr* promoter through unknown transcription factors based on the results of H3K4me3 ChIP-qPCR, and further researches are needed to identify which transcription factor ornithine affects and how ornithine regulates it.

## Conclusion

For the first time, we revealed that BBR can disrupt the spermatogenesis in mouse model. Mechanistically, ornithine produced by Muribaculaceae facilitate spermatogenesis by accelerating the transcription of *Ldlr*, which is essential for cholesterol transportation for testosterone biosynthesis and spermatogenesis. We integrated the results of the gut microbiota, fecal metabolome, and testicular transcriptome, which provides a method for exploring the function of TCM with low bioavailability. Considering the results, moreover, the fertility impact should not be ignored for male patients when BBR is used to treat diseases.

## Declaration

### Ethics approval

All animal protocols were approved by the Institutional Animal Care and Use Committee of the Wuhan University. All animal experiments were performed in accordance with the Guidelines for Animal Experiments.

### Data Availability

All data relevant to the study are included in the article or are uploaded as supplementary information. All raw sequences have been deposited in the NCBI Sequence Read Archive (SRA) database (https://www.ncbi.nlm.nih.gov/) with the following accession numbers: PRJNA978974 (RNA sequencing of testes), PRJNA898107 (16S-rRNA sequencing of BBR feces), and PRJNA989049 (16S-rRNA sequencing of PEG feces).

### Authors’ contributions

R.L., M.L., Y.H. and W.Q. designed the work. W.Q., Y.X. and J.L. conducted the experiments and drafted the manuscript. J.Y. offered advice. H.S., B.W and J.C. helped with the bioinformatics analysis. W.Q. and R.L. revised the manuscript. X.Y. and H.S. performed part of the experiment. All authors approved the final version of the manuscript.

## Acknowledgments

The authors thank Professor Huatao Chen and Master Yating Li from Northwest Agriculture & Forestry University for providing the TM3 cell line.

---

Fig. S1 The rarefaction curves of the bacterial community. **a-c** The rarefaction curves of Chao1 index (**a**), Shannon index (**b**) and Simpson index (**c**) were showed.

Fig. S2 PEG inhibited the *Muribaculaceae*. **a** The total fecal bacterial load and category were analyzed in untreated and PEG-treated groups. **b-d** The alpha diversity indices (Chao1 index (**b**), Shannon index (**c**) and Simpson index (**d**)) were analyzed. N=4 for each group. **e** The ratios of Firmicutes/Bacteroidota were analyzed at the phylum levels. N=4 for each group. **f** Taxonomic cladogram from linear discriminant analysis effect size (LEfSe) was showed. Dot size is proportional to the abundance of the taxon. N=4 for each group. Data are presented as mean±SEM. Differences without statistical significance were not labled, *p<0.05, **p<0.01, ***p <0.001.

Fig. S3 Phylogenetic tree shows the relation of *Muribaculaceae* isolates. Bar, 0.02 indicates nucleotide substitutions per position.

Fig. S4 Gene ontology (GO) analysis was showed with differentially expressed genes of testes from control and BBR-treated mice. **a** The cellular component involved in differentially expressed genes between untreated groups and BBR-treated groups was analyzed. **b** The molecular function involved in differentially expressed genes was analyzed.

Fig. S5 The decreased in mRNA levels of *Ldlr* was not due to its degradation, but to a decline in ornithine. **a** *Odc1* mRNA levels in the testes from control and BBR-treated mice were analyzed by RT-qPCR. N=4 for each group. **b** *Csde1* mRNA levels in the testes from control and BBR-treated mice were analyzed by RNA sequencing. N=2 for each for each group. Data are presented as mean±SEM. Differences without statistical significance were not labled, *p<0.05, **p<0.01, ***p <0.001.

## References

1. Sharlip ID, Jarow JP, Belker AM, Lipshultz LI, Sigman M, Thomas AJ, Schlegel PN, Howards SS, Nehra A, Damewood MD, et al: Best practice policies for male infertility. Fertil Steril 2002, 77:873–882.

2. Ding N, Zhang X, Zhang XD, Jing J, Liu SS, Mu YP, Peng LL, Yan YJ, Xiao GM, Bi XY, et al: Impairment of spermatogenesis and sperm motility by the high-fat diet-induced dysbiosis of gut microbes. Gut 2020, 69:1608–1619.

3. Chang Z, Qin W, Zheng H, Schegg K, Han L, Liu X, Wang Y, Wang Z, McSwiggin H, Peng H, et al: Triptonide is a reversible non-hormonal male contraceptive agent in mice and non-human primates. Nat Commun 2021, 12:1253.

4. Zhou DR, Zhou YC, Cui GH, Guo X, Qin J, Gui YT, Cai ZM: Gossypol repressed the gap junctional intercellular communication between Sertoli cells by decreasing the expression of Connexin43. Toxicol In Vitro 2008, 22:1719–1725.

5. Yan M, Wang L, Cheng CY: Testis Toxicants: Lesson from Traditional Chinese Medicine (TCM). Adv Exp Med Biol 2021, 1288:307–319.

6. Cui Y, Han J, Ren J, Chen H, Xu B, Song N, Li H, Liang A, Shen G: Untargeted LC-MS-based metabonomics revealed that aristolochic acid I induces testicular toxicity by inhibiting amino acids metabolism, glucose metabolism, beta- oxidation of fatty acids and the TCA cycle in male mice. Toxicol Appl Pharmacol 2019, 373:26–38.

7. Li L, Wang X, Sharvan R, Gao J, Qu S: Berberine could inhibit thyroid carcinoma cells by inducing mitochondrial apoptosis, G0/G1 cell cycle arrest and suppressing migration via PI3K-AKT and MAPK signaling pathways. Biomed Pharmacother 2017, 95:1225-1231.

8. Budeyri Gokgoz N, Avci FG, Yoneten KK, Alaybeyoglu B, Ozkirimli E, Sayar NA, Kazan D, Sariyar Akbulut B: **Response of Escherichia coli to Prolonged Berberine Exposure**. Microb Drug Resist 2017, 23:531–544.

9. Liu Y, Hua W, Li Y, Xian X, Zhao Z, Liu C, Zou J, Li J, Fang X, Zhu Y: Berberine suppresses colon cancer cell proliferation by inhibiting the SCAP/SREBP-1 signaling pathway-mediated lipogenesis. Biochem Pharmacol 2020, 174:113776.

10. Clemente JC, Ursell LK, Parfrey LW, Knight R: The impact of the gut microbiota on human health: an integrative view. Cell 2012, 148:1258–1270.

11. Li D, Liu R, Wang M, Peng R, Fu S, Fu A, Le J, Yao Q, Yuan T, Chi H, et al: 3beta- Hydroxysteroid dehydrogenase expressed by gut microbes degrades testosterone and is linked to depression in males. Cell Host Microbe 2022, 30:329–339 e325.

12. Huang G, Khan I, Li X, Chen L, Leong W, Ho LT, Hsiao WLW: Ginsenosides Rb3 and Rd reduce polyps formation while reinstate the dysbiotic gut microbiota and the intestinal microenvironment in Apc(Min/+) mice. Sci Rep 2017, 7:12552.

13. Wu J, Wei Z, Cheng P, Qian C, Xu F, Yang Y, Wang A, Chen W, Sun Z, Lu Y: Rhein modulates host purine metabolism in intestine through gut microbiota and ameliorates experimental colitis. Theranostics 2020, 10:10665–10679.

14. Hu X, Yu L, Li Y, Li X, Zhao Y, Xiong L, Ai J, Chen Q, Wang X, Chen X, et al: Piperine improves levodopa availability in the 6-OHDA-lesioned rat model of Parkinson’s disease by suppressing gut bacterial tyrosine decarboxylase. CNS Neurosci Ther 2023.

15. Li H, Li N, Lu Q, Yang J, Zhao J, Zhu Q, Yi S, Fu W, Luo T, Tang J, et al: Chronic alcohol-induced dysbiosis of the gut microbiota and gut metabolites impairs sperm quality in mice. Front Microbiol 2022, 13:1042923.

16. Tremellen K, Pearce K: Small intestinal bacterial overgrowth (SIBO) as a potential cause of impaired spermatogenesis. Gut 2020, 69:2058–2059.

17. Zhang P, Feng Y, Li L, Ge W, Yu S, Hao Y, Shen W, Han X, Ma D, Yin S, et al: Improvement in sperm quality and spermatogenesis following faecal microbiota transplantation from alginate oligosaccharide dosed mice. Gut 2021, 70:222–225.

18. Tropini C, Moss EL, Merrill BD, Ng KM, Higginbottom SK, Casavant EP, Gonzalez CG, Fremin B, Bouley DM, Elias JE, et al: Transient Osmotic Perturbation Causes Long-Term Alteration to the Gut Microbiota. Cell 2018, 173:1742–1754 e1717.

19. Su L, Mruk DD, Cheng CY: Drug transporters, the blood-testis barrier, and spermatogenesis. J Endocrinol 2011, 208:207–223.

20. Moon JM, Ratliff KM, Hagele AM, Stecker RA, Mumford PW, Kerksick CM: Absorption Kinetics of Berberine and Dihydroberberine and Their Impact on Glycemia: A Randomized, Controlled, Crossover Pilot Trial. Nutrients 2021, 14.

21. Lagkouvardos I, Lesker TR, Hitch TCA, Galvez EJC, Smit N, Neuhaus K, Wang J, Baines JF, Abt B, Stecher B, et al: Sequence and cultivation study of Muribaculaceae reveals novel species, host preference, and functional potential of this yet undescribed family. Microbiome 2019, 7:28.

22. Miyake S, Ding Y, Soh M, Low A, Seedorf H: Muribaculum gordoncarteri sp. nov., an anaerobic bacterium from the faeces of C57BL/6J mice. Int J Syst Evol Microbiol 2020, 70:4725-4729.

23. Gu J, Zhu N, Li HF, Zhao TJ, Zhang CJ, Liao DF, Qin L: Cholesterol homeostasis and cancer: a new perspective on the low-density lipoprotein receptor. Cell Oncol (Dordr) 2022, 45:709–728.

24. Gao Z, Liu S, Tan L, Gao X, Fan W, Ding C, Li M, Tang Z, Shi X, Luo Y, Song S: Testicular toxicity of bisphenol compounds: Homeostasis disruption of cholesterol/testosterone via PPARalpha activation. Sci Total Environ 2022, 836:155628.

25. Liu X, Wang C, Liu W, Li J, Li C, Kou X, Chen J, Zhao Y, Gao H, Wang H, et al: Distinct features of H3K4me3 and H3K27me3 chromatin domains in pre- implantation embryos. Nature 2016, 537:558–562.

26. Zhang W, Xia S, Xiao W, Song Y, Tang L, Cao M, Yang J, Wang S, Li Z, Xu C, et al: A single-cell transcriptomic landscape of mouse testicular aging. J Adv Res 2022.

27. Yuan L, Li Q, Bai D, Shang X, Hu F, Chen Z, An T, Chen Y, Zhang X: La(2)O(3) Nanoparticles Induce Reproductive Toxicity Mediated by the Nrf-2/ARE Signaling Pathway in Kunming Mice. Int J Nanomedicine 2020, 15:3415–3431.

28. Habtemariam S: Berberine pharmacology and the gut microbiota: A hidden therapeutic link. Pharmacol Res 2020, 155:104722.

29. Menees S, Saad R, Chey WD: **Agents that act luminally to treat diarrhoea and constipation**. Nat Rev Gastroenterol Hepatol 2012, 9:661–674.

30. Amin AH, Subbaiah TV, Abbasi KM: Berberine sulfate: antimicrobial activity, bioassay, and mode of action. Can J Microbiol 1969, 15:1067–1076.

31. Choudhry VP, Sabir M, Bhide VN: **Berberine in giardiasis**. Indian Pediatr 1972, 9:143–146.

32. Visconti A, Le Roy CI, Rosa F, Rossi N, Martin TC, Mohney RP, Li W, de Rinaldis E, Bell JT, Venter JC, et al: Interplay between the human gut microbiome and host metabolism. Nat Commun 2019, 10:4505.

33. Kenny DJ, Plichta DR, Shungin D, Koppel N, Hall AB, Fu B, Vasan RS, Shaw SY, Vlamakis H, Balskus EP, Xavier RJ: **Cholesterol Metabolism by Uncultured Human Gut Bacteria Influences Host Cholesterol Level**. Cell Host Microbe 2020, 28:245–257 e246.

34. Jiang F, Gao Y, Dong C, Xiong S: ODC1 inhibits the inflammatory response and ROS-induced apoptosis in macrophages. Biochem Biophys Res Commun 2018, 504:734–741.

35. Zirkin BR, Papadopoulos V: **Leydig cells: formation, function, and regulation**. Biol Reprod 2018, 99:101–111.

36. Hu J, Zhang Z, Shen WJ, Azhar S: Cellular cholesterol delivery, intracellular processing and utilization for biosynthesis of steroid hormones. Nutr Metab (Lond) 2010, 7:47.

37. Smith GA, Padmanabhan A, Lau BH, Pampana A, Li L, Lee CY, Pelonero A, Nishino T, Sadagopan N, Xia VQ, et al: Cold shock domain-containing protein E1 is a posttranscriptional regulator of the LDL receptor. Sci Transl Med 2022, 14:eabj8670.

