## Supplementary figures and images for "Berberine impairs spermatogenesis in mice by modulating ornithine metabolism via gut microbiota regulation"

### Fig S1.tif

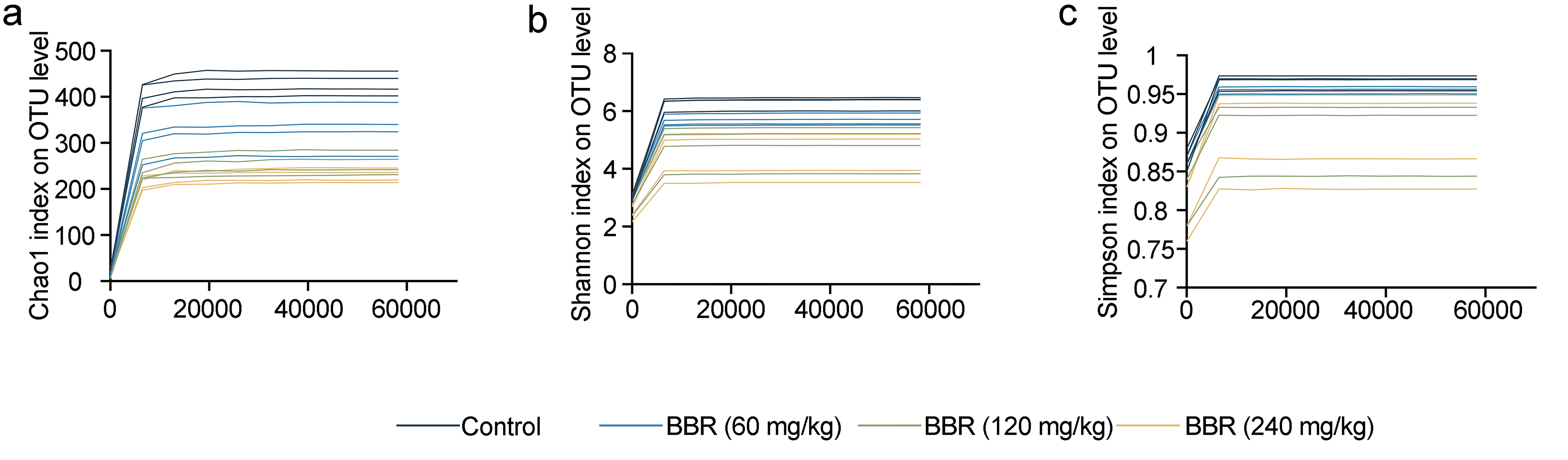

### Fig S2.tif

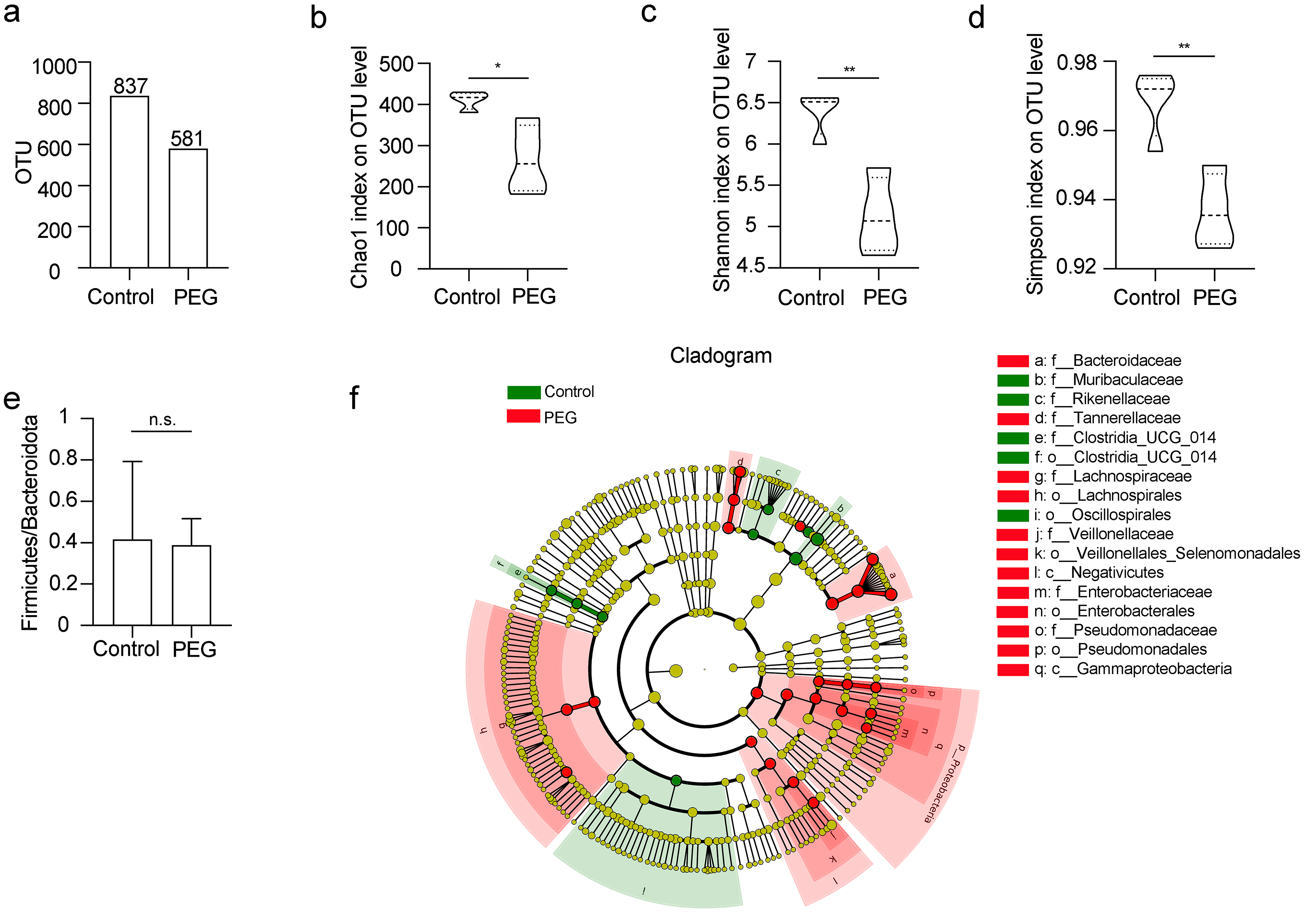

### Fig S3.tif

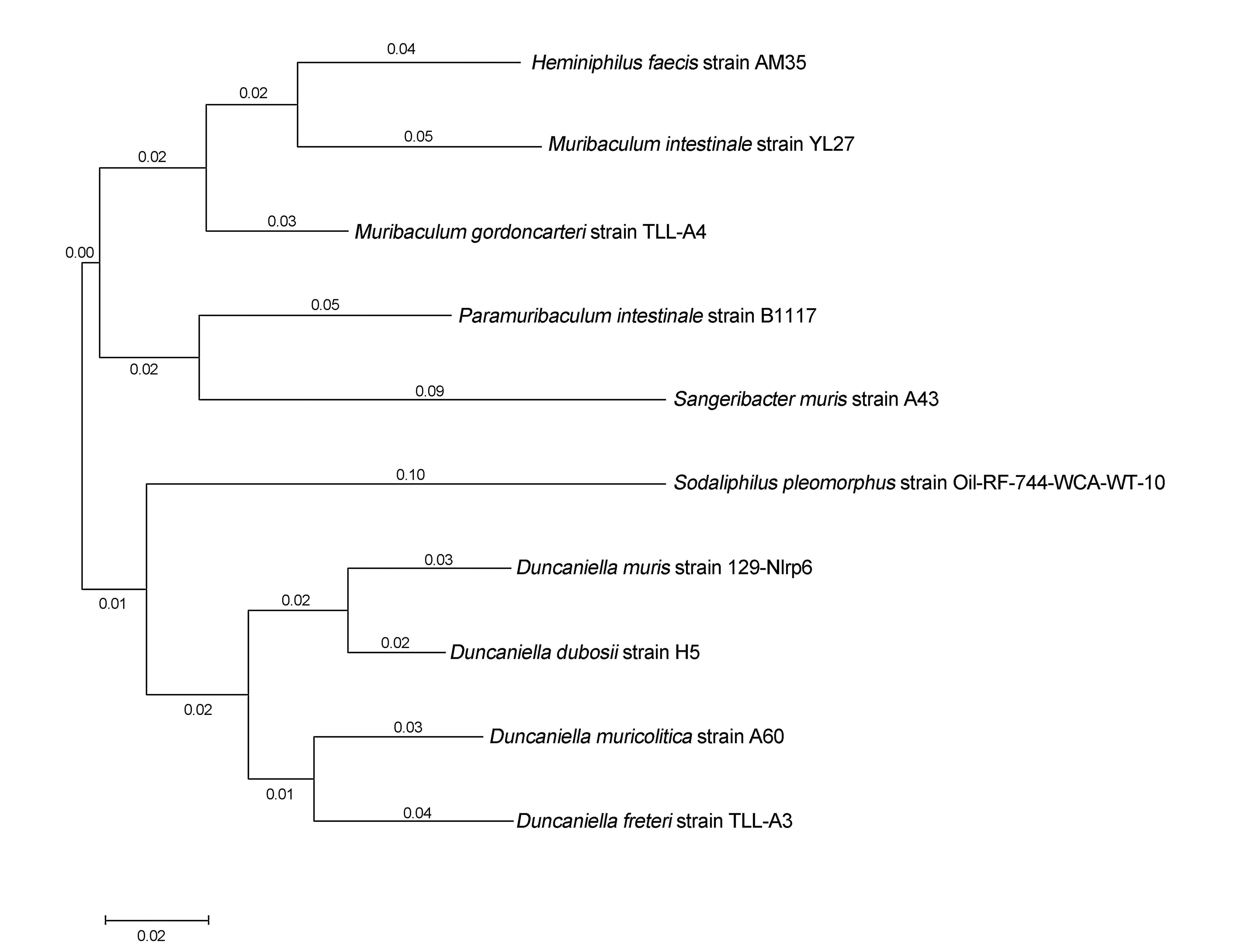

### Fig S4.tif

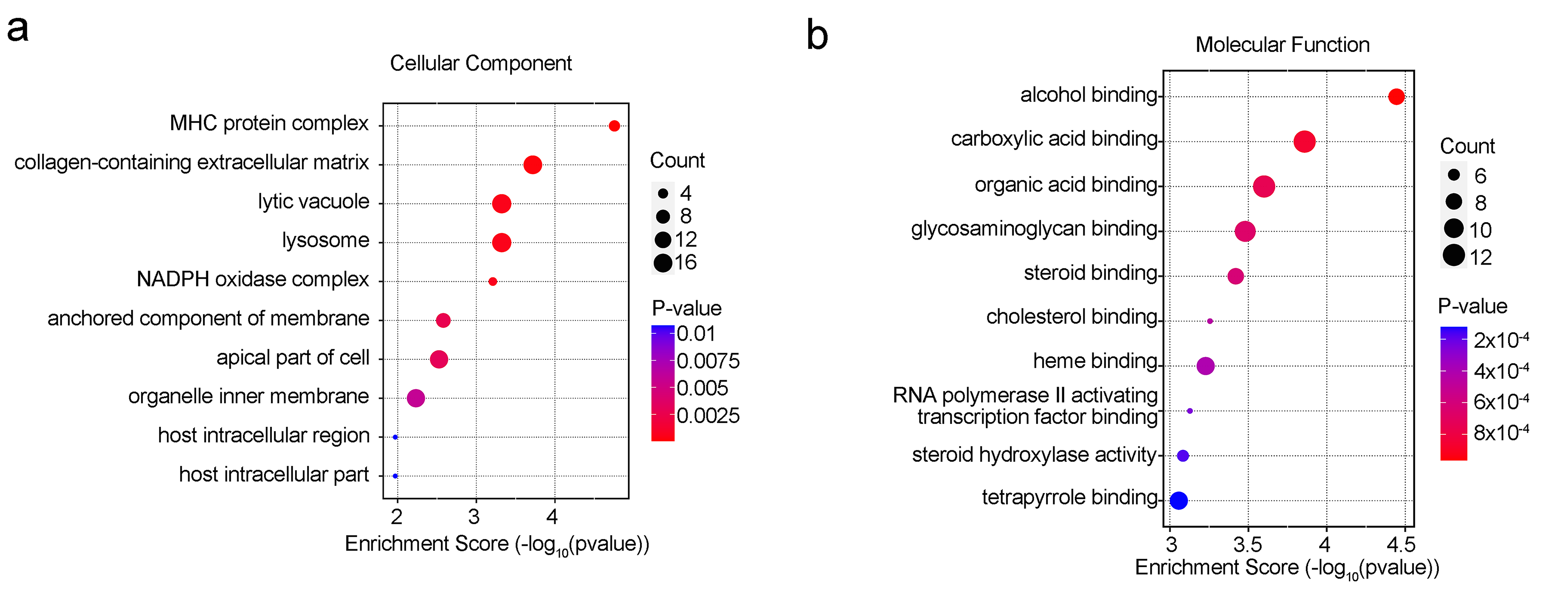

### Fig S5.tif

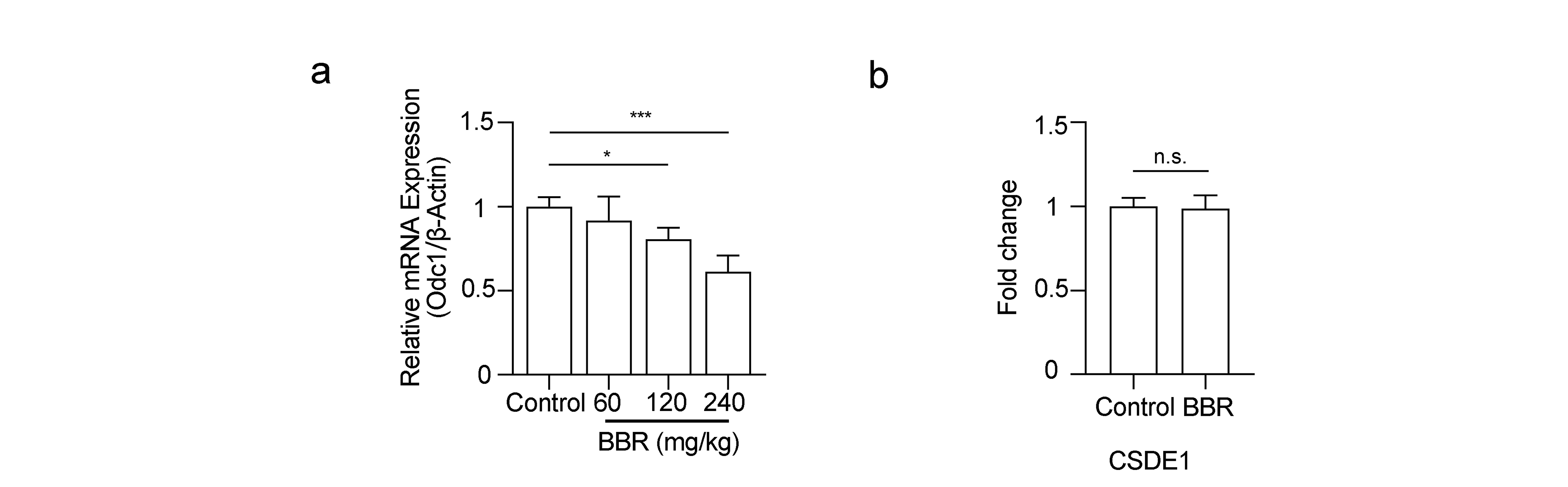
