## Supplementary tables for "Berberine impairs spermatogenesis in mice by modulating ornithine metabolism via gut microbiota regulation": Table S1.docx

**Table S1. Antibodies used for immunofluorescence staining.**

| **Antibody** | **Cat. #** | **Source** | **Company** |
| --- | --- | --- | --- |
| Anti-ZO-1 antibodies | 21773-1-AP | Rabbit | Proteintech |
| CoraLite488 goat anti-rabbit IgG | SA00013-2 | Goat | Proteintech |
