## Supplementary tables for "Berberine impairs spermatogenesis in mice by modulating ornithine metabolism via gut microbiota regulation": Table S2.docx

**Table S2. Primer sequences used for qRT-PCR.**

| **Gene** | **Sequence 5’-3’** | **Amplicon (bp)** |
| --- | --- | --- |
| *Plzf* | F: CTGGACAGTTTGCGACTGAG | 123bp |
|  | R: GTCTGTGTGTCTCCA |  |
| *Hsd3b1* | F: TGTCACAGGTGTCATTCCCA | 113bp |
|  | R: AGCTGCAGAAGATGAAGGC |  |
| *Sycp3* | F: GGACAGCGACAGCTCAC | 100bp |
|  | R: ATCAACCAAAGGTGGCTTCC |  |
| *Sox9* | F: AGCTGGCAAAGTTGATCTG | 136bp |
|  | R: GACGTCGAAGGTCTCAATGTTGG |  |
| *Cyp11a1* | F: AGGAGACACTGAGACTCCAC | 132bp |
|  | R: GATCTCGACCCATGGCAAAG |  |
| *Cyp17a1* | F: AATGAATGGGACCAGCCAGA | 93bp |
|  | R: GGGCAAATAACTGGGTGTGG |  |
| *StAR* | F: CAGGAAGGCTGGAAGAAGGA | 199bp |
|  | R: TCTGCAGGACCTTGATCTCC |  |
| *β-actin* | F: GAGACCTTCAACACCCCAGCC | 362bp |
|  | R: CCGTCAGGCAGCTCATAGCTC |  |
| *Ldlr* | F：TCCACTGTGGTAGCAGTGAG | 146bp |
|  | F：GTGAATGCAGGAGCCATCTG |  |
| *Ldlr-pro* | F: TGGAAACCTCGCCCCTAGTA | 140bp |
|  | R: AATGTTCCCGCTGCAAACAC |  |
| *AR* | F: GGCGGTCCTTCACTAATGTCAACT | 79bp |
|  | R: TGTGCATGCGGTACTCATTG |  |
